# Characterizing *Xenopus tropicalis* endurance capacities with multilevel transcriptomics

**DOI:** 10.1101/091280

**Authors:** Adam J. Richards, Anthony Herrel, Mathieu Videlier, Konrad Paszkiewicz, Nicolas Pollet, Camille Bonneaud

## Abstract

Vertebrate endurance capacity is a phenotype with considerable genetic heterogeneity. RNA-Seq technologies are an ideal tool to investigate the involved genes and processes, but several challenges exist when the phenotype of interest has a complex genetic background. Difficulties manifest at the level of results interpretation because commonly used statistical methods are designed to identify strongly associated genes. If an observed phenotype can be achieved though multiple distinct genetic mechanisms then typical gene-centric methods come with the attached risk that signal may be lost or misconstrued.

Gene set analysis (GSA) methods are now widely accepted as a means to address some of the shortcomings of gene-by-gene analysis methods. We carry out both gene level and gene set level analyses on *Xenopus tropicalis* to identify the genetic factors that contribute to endurance heterogeneity. A typical workflow might consider gene level and pathway level analyses, but in this work we propose an additional focus at the intermediate level of functional modules. We generate functional modules for GSA testing in order to be explicit in how ontology information is used with respect to the functional genomics of *Xenopus*. Additionally, we make use of multiple assemblies to corroborate implicated genes and processes.

We identified 42 core genes, 10 functional modules, and 14 pathways based on gene expression differences between endurant and non-endurant frogs. The majority of the genes and processes are readily associated with muscle contraction or catabolism. A substantial number of these genes are involved in lipid metabolic processes, suggesting an important role in frog endurance heterogeneity. Unsurprisingly, many of the gene expression differences between endurant and non-endurant frogs can be distilled down to the capacity to utilize substrate for energy, but at the individual level frogs appear to make use of diverse machinery to achieve these differences.

## 1 Background

While RNA-Seq technologies offer tremendous promise for the identification of genes and processes underlying complex phenotypes, the biological interpretation of results remains challenging. This can come from an incomplete picture of the underlying biology, the use of inferred annotations or it may be an artifact that arises because gene-centric methods are not well-suited for genetically diverse phenotypes. The results of gene expression studies invariably take the form of one or more gene lists and bridging the gap between these lists and the relevant biology can be a major hurdle (Huang et al., 2009; Richards et al., 2010). The problems that come with interpreting these lists are all the more exacerbated in non-model organisms, which for the most part, face the added difficulty of a poorly annotated genome. Gene Set Analysis (GSA) methods can help alleviate the issues that come with the use of gene-centric methods, by focusing on pathways or processes instead of the genes themselves (Mootha et al., 2003; Subramanian et al., 2005). These methods make use of gene sets determined *a priori* and they are most commonly applied to pathway analysis. To aid results navigation and to address the issues arising from inferred annotations we consider, in addition to the gene and pathway level analysis, the intermediate level of functional modules.

Functional modules have been identified from protein-protein interaction networks (Dittrich et al., 2008) as well as from expression data (Tully et al., 2014) and for some time now there has been a general consensus that functional modules are important building blocks in biological systems (Spirin and Mirny, 2003; Tanay et al., 2004). Functional modules have been used to understand biological systems by analyzing various representations of the systems themselves, but the hypothesis testing framework GSA has only rarely (if ever) been explicitly used in this context. The methods presented here make use of functional modules determined *a priori* and they explicitly consider functional annotations that are relevant to frog species. This is an important and novel aspect of this study because, it is our understanding that there are no methods in the literature that generate species-specific functional modules for GSA testing.

This study presents a novel transcriptome of a wild amphibian, the western clawed frog *Xenopus tropicalis* using a multi-level analysis approach. We characterize the genetic components that contribute to the spectrum of frog locomotor capacity, a fundamental aspect of animal movement and population dynamics (Morales et al., 2010). Endurance is a complex trait that is likely to be under strong selection in natural populations in response to human activities and widespread environmental change. Direct (e.g. habitat destruction) and indirect habitat alterations (e.g. climate change, habitat fragmentation) should increase risks of local population extinction by reducing the number of suitable habitats and their connectivity, such that population persistence through the colonization of novel habitats will, in part, depend on the ability to disperse over potentially long distances. Accurate ecological forecasting of population dynamics and persistence are, however, limited by our power to predict whether and how fast organisms will respond to selection on dispersal and mobility. This, in turn, will require an improved understanding of the molecular underpinnings of variation in mobility in natural populations. While large scale genetics studies on locomotor capacity have been carried out for laboratory populations of Drosophila melanogaster (Jordan et al., 2007), our understanding of the genetic basis of locomotion and mobility in wild vertebrates, including amphibians,remains largely incomplete.

Amphibians are currently among the most endangered vertebrate taxa, with over a third of species threatened globally (Wake and Vredenburg, 2008; Blaustein et al., 2010). Habitat fragmentation and loss are a major cause of amphibian decline and they impact 90% of species threatened (Stuart et al., 2004). In particular, the mature tropical forests containing the majority of all amphibians are not only rapidly shrinking as a result of logging, but are also predicted to experience extreme climatic stress in the coming decades that will lead to their aridification and fragmentation (Beaumont et al., 2011; Zelazowski et al., 2011). Climate change is similarly forecasted to impact amphibians, not only because they have evolved temperature optima that closely match environmental temperatures (Angilletta and Frazier., 2010), but also because adults generally show limited acclimation capabilities (Wilson et al., 2000). As a result, we expect amphibians experiencing such dramatic environmental change to be under strong selection for dispersal to more suitable habitats. *X. tropicalis* has a natural distribution restricted to Western Africa, is largely aquatic and confined to acidic rainforests pools (Rödel, 2000). In this species, terrestrial locomotion plays an important role in overland dispersal during periods of heavy rainfall (Rödel, 2000).

Over the last 10 years, *X. tropicalis* has emerged as a model species in genomics, due to its embryological and developmental similarities with the classic model species, *X. laevis* (e.g. Gurdon et al., 1958), but it is easier to work with due to a shorter generation time and smaller diploid genome that qualifies as relatively simple among amphibians (Hirsch et al., 2002). Research in *X. tropicalis* genomics began with the production of cDNA clones for expressed sequence tag (EST) projects (Gilchrist et al., 2004; Morin et al., 2006; Fierro et al., 2007) and reached an important milestone with genome publication (Hellsten et al., 2010). As a result, a number of *Xenopus*-centric resources have emerged (Sczyrba et al., 2005; Bowes et al., 2008), which the repository Xenbase (http://www.xenbase.org) serves to centralize (James-Zorn et al., 2013). Unfortunately, the *X. tropicalis* genome has only sparse functional annotation. Although efforts exist (Schmid and Blaxter, 2008) to infer annotation in non-model species from sequence similarity there is still a significant knowledge gap between most organisms and the few most widely studied ones.

The inference of annotation information in non-model species is by no means novel and complex methods have existed for several decades so a review of the numerous approaches is well beyond the scope of this work. Regardless of the method employed it is important that the organisms used to carry out inference be those that make the most sense within the context of the experimental system. In this work we analyzed whole-genome RNA expression data from endurant and non-endurant *X. tropicalis* individuals with a focus on combining annotation information from *X. tropicalis* and *X. laevis*. Principally this was done through the generation of functional modules that were tested using the GSA framework (Efron and Tibshirani, 2007). GSA has the advantage of taking into account the modular architecture of transcriptome organizations (Tanay et al., 2004), which helps facilitate the identification of small coordinated changes in expression that may go undetected with gene-centric statistics (Mootha et al., 2003; Subramanian et al., 2005).

Gene expression studies make use of accumulated biological knowledge, not just at the level of genome annotation (coding region mapping, non-coding elements etc.), but also at the level of functional, structural and pathway annotation. In the functional regard, the Gene Ontology (GO) has become an immensely useful tool in genomics (Ashburner et al., 2000; Consortium, 2015). Through the GO individual genes may be mapped to appropriate annotations to provide important context irregardless of the study system. Unfortunately, even if we consider electronically inferred annotations *X. tropicalis* is only sparsely annotated compared to other organisms in the GO database. For non-model organisms this is a common scenario, but some effort has been made to help researchers infer annotations from sequence data (Schmid and Blaxter, 2008). In this work we combine *X. tropicalis* and *X. laevis* annotation data to create composite functional modules for testing within the framework of gene set analysis (GSA) (Subramanian et al., 2005; Efron and Tibshirani, 2007).

In order to provide perspective on the genes and processes associate with *X. tropicalis* endurance heterogeneity we analyzed expression differences using three distinct transcriptome assemblies. Specifically, we assembled the reads as *de novo*, genome guided and reference, making use of the *X. tropicalis* genome for the latter two. Reference assemblies tend to produce longer and higher quality transcripts than *de novo* ones, but *de novo* methods have the advantage of identifying novel or poorly annotated gene regions (Tan et al., 2013; Marchant et al., 2015). Using multiple assemblies, at least for *X. tropicalis*, helps control for the inherent biases that come with the different transcriptome assembly procedures. Comparing and contrasting the results at the individual gene, functional module, and pathway level, across the three assemblies, provided a reasonable way to prioritize results for further experimental and scientific inquiry.

## 2 Results

### 2.1 RNA-Seq assemblies

*X. tropicalis* has a relatively small haploid genome with about 1.7 Gbp making up 10 chromosomes (Tymowska, 1973). The genome was sequenced and assembled such that 97.6% of the known coding genes were contained within the reported scaffolds (Hellsten et al., 2010). The reads in this study, obtained from Illumina sequencing, were assembled using *de novo* (DN) and genome-guided (GG) *de novo* Trinity (Grabherr et al., 2011). Additionally, the reads were aligned and mapped directly to the most recent genome build, using STAR, thereby providing a reference assembly (RF) (Dobin et al., 2013). The *X. tropicalis* reference genome (JGI assembly v7.2) contained 7730 scaffolds (James-Zorn et al., 2013). The DN, GG and RF assemblies consisted of 218,541 52,543 and 28,171 transcripts respectively. Trinity creates and makes use of a composite assembly to align and map sample reads, where STAR, aligns the reads directly to the reference genome. As a consequence, the reference assembly is reported in terms of isoforms, whereas Trinity assemblies also contain an estimate of the gene level counts (DN = 163,981 and GG = 41,256). As is typical with *de novo* assemblies, the estimated number of genes is generally higher than the expected gene count. Because we compare amongst assemblies to prioritize genes and processes, results are reported using an isoform treatment of the data.

Tan and colleagues recently published the developmental transcriptome of *X. tropicalis* and their reference assembly covered ∼97.3% (8211/8437) of the the known annotated RefSeq genes (Tan et al., 2013). Although we are mapping to a more recent RefSeq database, it is still interesting to report that our reference assembly covered ∼88.9% (8286/9319) of known annotated genes. The *de novo* (7086/9319) and genome-guided (5906/9319) assemblies represented fewer annotated genes, which results in lower reference coverage. It is reasonable to expect a lower percentage of genes expressed for a phenotype of locomotor activity compared to the entire developmental process and this should at least partly account for the observed differences. If we use the SwissProt portion of the UniProt database in addition to the genome build, we observe that the transcripts from *de novo* assemblies map to more proteins (Consortium, 2014). Specifically, we see that for the DN, GG, and RF assemblies there are 20,359, 14,442, 16,047 unique genes after filtering away genes that do not map to a protein (E-value threshold ≤ 0.0001). Many mapped genes will turn out to be homologs from different species, ultimately reducing the realized total genes, but the higher number for the *de novo* assembly attests to the discoverability of *de novo* approaches.

### 2.2 Transcript level expression analysis

The first step in our tri-level approach to gene expression analysis was at the level of individual genes. Differentially expressed genes were determined with respect to `endurant` and `non-endurant` classifications. Samples were assigned a class based on a Gaussian mixture model (see Fig. 1A-C). The mixture model incorporated four criteria that summarized one or more performance tasks: maximum(stamina), maximum(swim speed), maximum(acceleration) and maximum(distance jumped) (see Methods). The maximum distance jumped is the principal discriminator between classes under this clustering and along with any of the other features the samples may readily be separated. Although, sample D is classified as `endurant` it is the one example of a frog with modest performances. A genetic distinction between phenotypes is supported by the degree of separation among `endurant` and `non-endurant` samples as shown in the column-wise dendrograms of Figure 1. The hierarchical clusterings are shown along with a corresponding heatmap of normalized counts (see panels D and E). The differential expression analysis and count transformation was carried out with DESeq2 Love et al. (2014).

**Figure 1:**
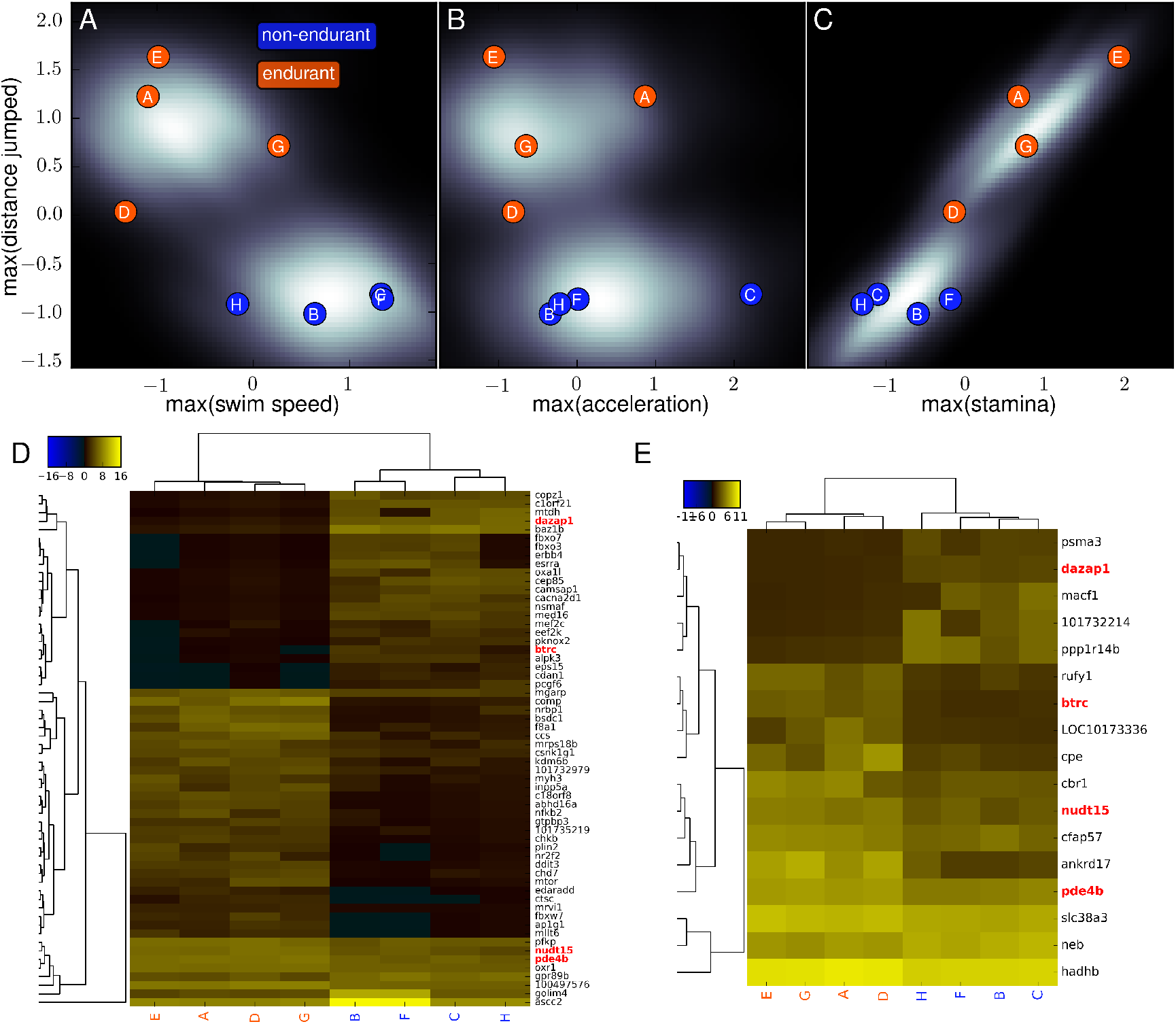
Transcript expression analysis. **A-C.** The performances of individual samples are overlaid on the modes of a Gaussian mixture model where the features are 4 different performance criteria. The samples show reasonable separation as `endurant` and `non-endurant` individuals. All features were standardized to have zero mean and unit variance. **D.** Heatmap representation of the regularized log transformed counts for the *de novo* assembly. To aid visualization, only one transcript per gene is shown and only a subset of the original transcripts are shown (50/97). **E.** For the genome-guided assembly, all transcripts that mapped to a coding gene are shown as transformed counts. For both heatmaps, hierarchical clustering was performed on the genes as well as the samples and all genes shown had significance levels ≤ 0.05. Note that over and under expression of transcripts in these heatmap representations is relative to the whole transcriptomes to allow for comparison across assemblies. The two assemblies share five genes (marked in green) one of which is not shown (*slc38a3*) in D because of the visualization truncation. Additionally, the color encoding for class labels in A-C is reiterated in the column labels of D and E.

To identify the individual genes that contribute the most to the phenotypic heterogeneity a series of filters were imposed on the transcripts. Transcripts were removed if there were no counts or if they did not pass the dispersion outlier criteria implemented as part of DESeq2 (Cook, 1977; Love et al., 2014). Finally, a permissive and non-permissive adjusted *p*-value threshold (≤ 0.5 and ≤ 0.05) filter was imposed. Using the permissive criteria, the DN, GG, and RF transcriptomes yielded 533, 89 and 46 transcripts respectively. The non-permissive criteria resulted in 97, 17 and 4 transcripts for the same assemblies. The heatmaps in Figure 1 correspond to the non-permissive criteria. The 4 significant transcripts from the RF assembly were aligned to the gene symbols *dgat2, prdx5, pvalb* and *LOC1004 87011*. Transcripts that correspond to *pvalb* are significantly differentially expressed (adj *p*-value ≤ 0.05) in the DN and RF assemblies. No genes are shared between the RF and GG assemblies at the non-permissive level of thresholding; however, the GG and DN assemblies share five genes at this cutoff: *slc38a3, nudt15, dazap1, pde4b*, and *btrc*.

While the heatmaps in Figure 1 serve to represent the individual heterogeneity in expression with respect to the most important genes, the volcano plots provide a perspective that averages over the individuals while highlighting the directionality of the fold change (see Fig. 2). The transformed values in the three subplots enable a direct comparison of expression values between assemblies. The DN assembly has the most significant genes and the most extreme values of significance (larger values on the y-axis indicate smaller *p*-values). As we expected the genome-guided assembly, a hybrid approach between the RF and DN assemblies, produces a distribution of results that appear to be a compromise between the extremes of the other assemblies. The top eight transcripts for each assembly are shown and *dazap1* along with *pde4b* are both present in two assemblies.

**Figure 2:**
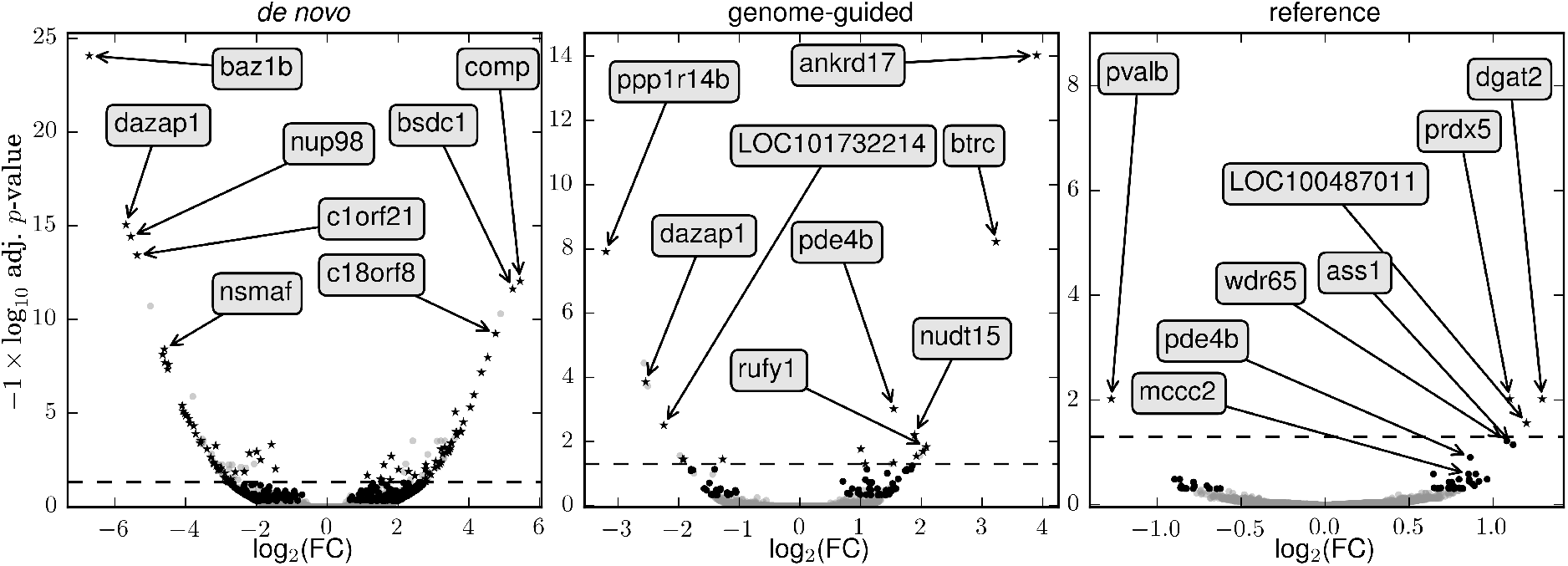
Transcript expression by assembly. The log_2_ fold changes are plotted against log_10_ transformed adjusted *p*-values for the three transcriptome assemblies. The directionality of change with respect to non-endurant frogs is indicated by on the x-axis. For each ‘volcano’ plot the top eight genes are annotated with arrows. The dashed line corresponds to the non-permissive threshold (adj. *p*-value ≤ 0.05). Transcripts that passed outlier thresholding, but had adjusted *p*-values ≥ 0.5 are plotted in grey. Transcripts that are ≤ 0.5 are shown in black and transcripts with adj. *p*-values ≤ 0.05 are shown as stars. The grey markers in the statistically significant region are alternative transcripts of those already marked with a star.

The most significant gene in the DN assembly *baz1b* is downregulated (negative fold change) in endurant frogs (see Fig.1D). These results provide evidence that the use of multiple assemblies gives a more complete picture of endurance heterogeneity in *X. tropicalis*.

### 2.3 Comparing transcriptomes to identify core genes

There are a number of promising genes that have been implicated by the gene level analyses (e.g. *dgat2* and *baz1b*), but there are still over 500 transcripts that pass the permissive threshold and for some applications it is necessary to further reduce the list. The non-permissive filter is very stringent and has a low tolerance for expression differences within a phenotypic class so a conservative *p*-value threshold is not an ideal method to filter for this study. The rationale is that performance heterogeneity is a complex phenotype and we do not expect all individuals to use identical mechanisms to achieve the observed phenotypic differences. Moreover, there is a fairly conservative logic in setting a permissive threshold, then averaging over the results from distinct transcriptomes. In essence each assembly is a different model of the data and we propose in this work to identify core genes by looking at the genes that lie at the intersection of any two assemblies. The transcripts and intersection regions are shown in Figure 3A. There are nine genes that are common to all assemblies are: *dgat2, wdr65, lpl, cpe, c14orf159, prdx5, bdh1, hadhb*, and *mccc2*. There are 42 genes that lie at the intersection of two or more assemblies. The 42 genes are laid out in an oval as yellow nodes in a network in Figure 3. Of the 42 genes 18 have at least one assembly that has an adjusted *p*-value ≤ 0.05 (shown as larger nodes).

**Figure 3:**
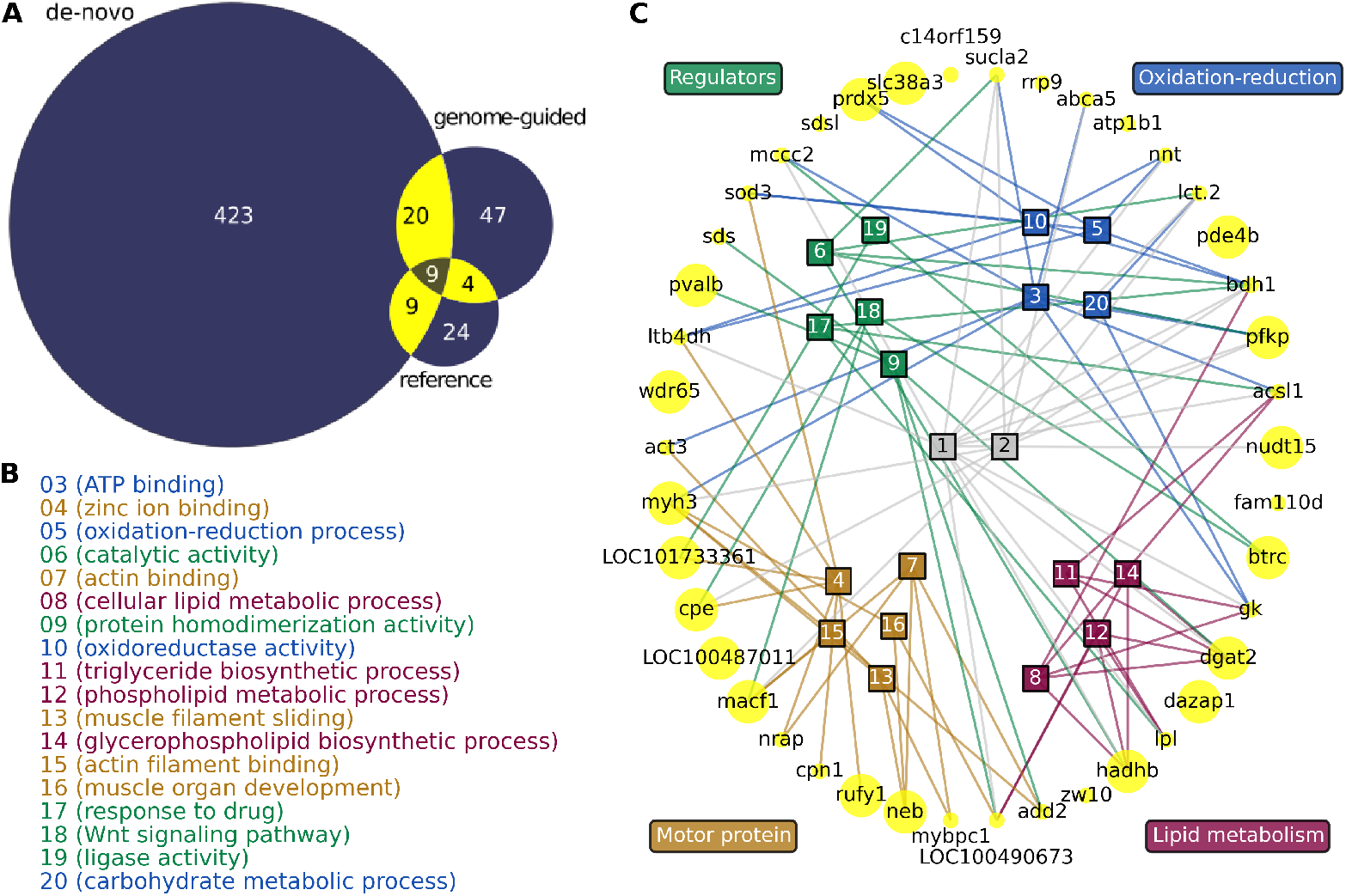
Genes at the intersection of multiple assemblies. **A** The total number of transcripts that are under the permissive (*p*-value ≤ 0.5) threshold are shown for each assembly. The number of transcripts that map to overlapping genes are shown at the intersection of the sets. **B** The legend for the network provides ontology term definations color coded by theme. **C** The 42 transcripts taken from the yellow region of the venn diagram are shown in a network using a circular layout. The corresponding gene symbols are used to label the nodes with larger sized ones indicating that at least one assembly had an adjusted *p*-value ≤ 0.05. The edges between ontology terms and genes are annotations and only the statistically significant ontology terms with at least three edges are shown. The ontology terms are broadly classified by color indicated theme as a visual aid. The terms 01 and 02 corresponding to *small molecule metabolic process* and *metabolic process* were left uncategorized because they are very general.

The 42 core genes have been sorted according to GO theme using the network in Figure 3C. The edges are annotations inferred from analysis of sequence homology with *Homo sapiens* for the biological process and molecular function aspects of the GO. The categories that are indicated by coloring are meant to loosely group the terms in order to better conceptualized the relationships among the genes. For example, the term *carbohydrate metabolic process* is not directly involved in redox reactions, but it is an important precursor. The category ‘regulation’ is meant to encompass regulation at the RNA, protein and metabolite levels and may be mediated via epigenetics, pathway modulation and a number of other mechanisms (see disucssion). The classification of ontology terms is mostly evident however a few terms need explanation. Protein dimerization is associated with many of these forms of regulation (Marianayagam et al., 2004). Regulation of zinc in skeletal muscle has been shown to be important for muscle contraction in the northern leopard frog (*Rana pipiens*) (Isaacson and Sandow, 1963). There are other connections among the genes, but we only show the terms that were significant (*p*-value ≤ 0.05) after testing for enrichment.

### 2.4 Functional module analysis

Because of corrections for multiple testing and filtering the analysis at the level of individual transcripts is constrained when trying to account for genetic heterogeneity. GO enrichment at this level reveals the processes that are affect by the most dramatically differentially expressed genes. There are however processes that may be affected in less extreme, but nonetheless biologically meaningful ways. To scan for processes that are more subtlety associated with our phenotype we generated functional modules specific to the genus *Xenopus*. The functional modules or gene sets for this account for the annotation information for *X. laevis* (NCBI Taxonomy ID: 8355) and *X. (Silurana) tropicalis* (NCBI Taxonomy ID: 8364). Because of the sparsity of annotation information we discuss here results with Inferred from Electronic Annotation (IEA) annotations. Both sets of results are described in more detail in the supplemental results. The bulk of annotations in the GO fall under this evidence category and the quality of these annotations has significantly improved in recent years (Skunca et al., 2012). Functional module collections were generated for both the molecular function and biological process aspects of the GO. We considered the 24817 NCBI gene identifiers that were specific to *X. tropicalis* and using annotation information from both species of frog we generated 157 functional modules for molecular function and 54 for biological process (see Methods). These modules were tested for enrichment with respect to the two phenotypes using the R package GSA (Efron and Tibshirani, 2007). The results are displayed in Figure 4.

**Figure 4:**
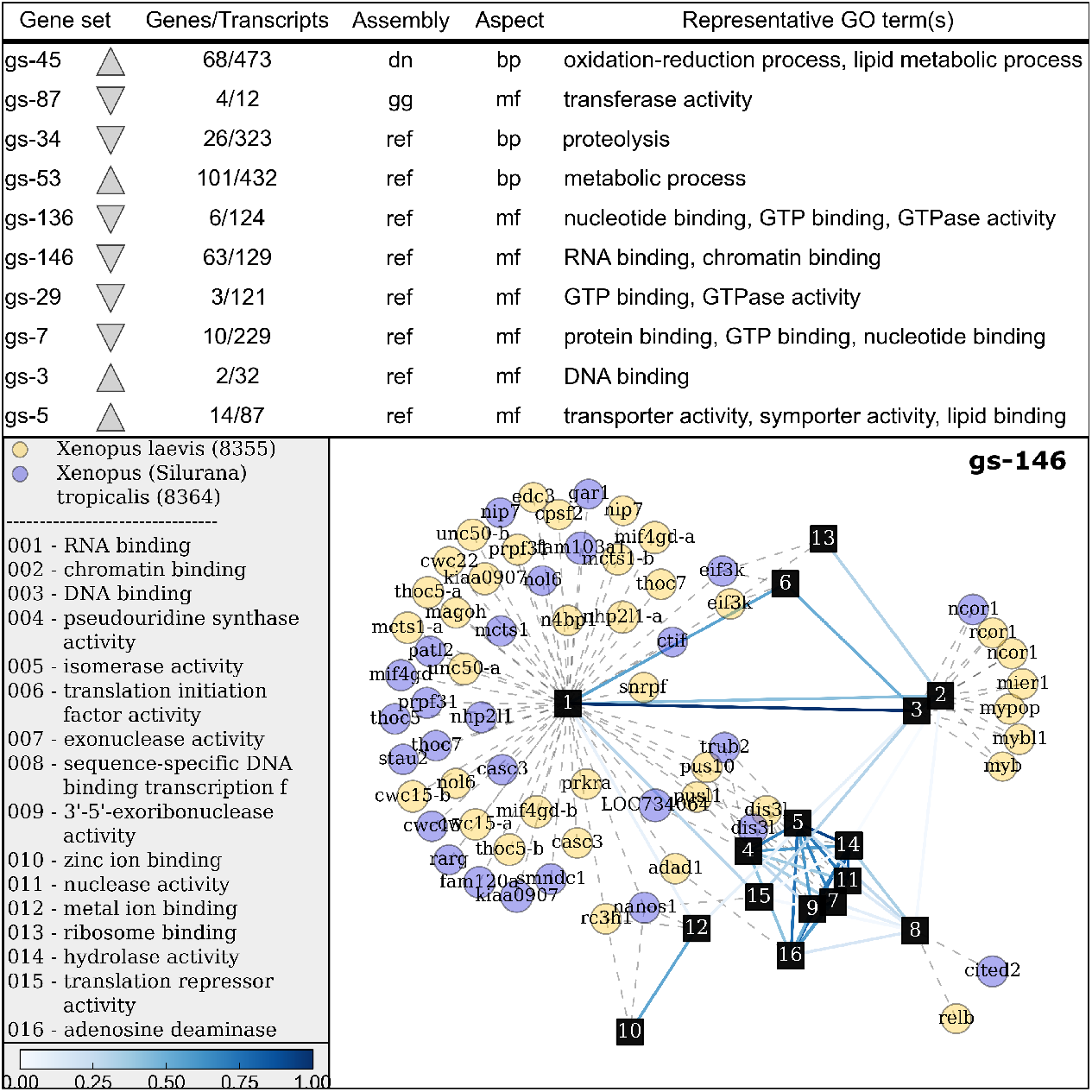
Gene set analysis. A total of 211 functional modules were tested for expression differences between endurant and non-endurant frogs using gene set analysis. The functional modules that were enriched with expression differences are shown in the top panel with their directionality indicated by the triangle. All ten gene sets had adjusted *p*-values ≤ 0.05. Additional information about the gene sets including the gene to transcript ratio are provided in the table. The bottom panel shows an example of on of the ten functional modules represented as a network. Annotations are designated with dashed line from terms (square nodes) to genes (circles). The color of the gene nodes indicate the species of *Xenopus*. The functional coherence of the genes in ‘gs-146’ is indicated by the number of interconnected edges among terms. The module is a group of genes likely connected to muscle contraction through zinc ion binding with regulation occurring via RNA, chromatin, or DNA binding.

The transcripts available for significance testing with GSA are those that can be matched with coding-genes because the annotations used in this study were specific to the GO. The central idea behind testing genes at the group level is that biological processes are more often than not carried out by multiple genes and accordingly we expect some level of coordination among genes that make up the process (Mootha et al., 2003). We investigated if coordination had occurred in the 211 generated functional modules. Many of the groups that are significantly associated with differences in phenotype essentially confirm the results of the individual transcript analysis, but a number of genes and processes with less dramatic expression differences were revealed. Many of the new processes are (shown top panel in Figure 4 and in the supplemental results) appear to be regulatory processes apparently intervening by binding be it DNA, RNA, chromatin or protein. Interestingly, GTP binding and GTPase has be implicated and there is evidence that GTP activity plays a role in regulation of smooth muscle contraction (Puetz et al., 2009).

Because the functional modules were generated using an unsupervised approach statistical significance does not necessarily imply biological significance and this is particularly true in the RNA-Seq setting. Functional modules were generated in gene space independent of the transcriptomes and in order to carry out the significance testing they were mapped to transcript space. Usually, there are multiple transcripts per gene as indicated by the top panel in Figure 4. Significant modules like gs-87, gs-136, gs-29, gs-7 and gs-3 all have a very small gene to transcript ratios and the significance in these gene sets may be an artifact of the over-representation of a given gene. It is difficult to completely rule out the possibility of biological significance in these cases because the significance may reveal an alternative preference for isoforms, which can be important (Wang et al., 2008). On the other hand gs-53, although significant has only a single term ‘metabolic process’ and in these cases there is simply not enough detail to assess biological significance. The functional modules with the most biological relevance are those with multiple genes and multiple (highly connected) terms. The module gs-146 (shown in Figure 4) is potentially reveals a bridge between regulatory activity and muscle contraction and the module gs-45 contains a number of core genes potentially indicating less dramatically expressed contributers to phenotypic heterogeneity.

## 3 Pathway Analysis

Functional modules are an intermediate level, falling between individual genes and biochemical pathways, with which we conceptualize biological activity. In GSEA the most commonly studied level is that of biological pathways (Tian et al., 2005) and when there is enough functional information available they may be thought of as the product of multiple functional module building blocks (Richards et al., 2012). We tested all downloadable KEGG pathways with the R package GSA as was done with the functional modules. The pathways and their directionality with respect to the non-endurant phenotype are shown in Table 1. Nearly every significant pathway (adjusted *p*-value ≤ 0.00001) is either directly or indirectly related to energy production. The pentose phospate pathway is a primarily anabolic form of metabolism and like glycerolipid metabolism it is appears to be restricted in endurant organisms. There are six pathways in all that are down-regulated under the endurant phenotype in contrast to ten pathways that are up-regulated. Four out of the ten up-regulated pathways were significant in two or more transcriptome assemblies and only a single pathway, propanoate metabolism, was significant in all three assemblies.

**Table 1:** The pathways that were significantly enriched with differentially expressed transcripts. Each pathways listed is associated with a difference in gene expression, between non-endurant and endurant frogs. A negatively (neg) enriched pathway indicates that the endurant transcripts were significantly down-regulated compared to their endurant counterparts. The third column indicated the assemblies that produced significant adjusted *p*-values (≤ 0.00001). The abbreviations correspond to the *de novo*,(dn), genome-guided (gg) and reference (rf) transcriptome assemblies.

|  |  |  |  |
| --- | --- | --- | --- |
| gs-00030 | Pentose phosphate pathway | dn | neg |
| gs-00561 | Glycerolipid metabolism | gg | neg |
| gs-03460 | Fanconi anemia pathway | dn | neg |
| gs-00040 | Pentose and glucuronate interconversions | gg | neg |
| gs-00510 | N-Glycan biosynthesis | gg | neg |
| gs-04140 | Regulation of autophagy | dn | neg |
| gs-00310 | Lysine degradation | dn,gg | pos |
| gs-00072 | Synthesis and degradation of ketone bodies | dn | pos |
| gs-00630 | Glyoxylate and dicarboxylate metabolism | dn | pos |
| gs-00071 | Fatty acid degradation | rf | pos |
| gs-00280 | Valine, leucine and isoleucine degradation | dn,gg | pos |
| gs-00900 | Terpenoid backbone biosynthesis | dn | pos |
| gs-00380 | Tryptophan metabolism | dn | pos |
| gs-00620 | Pyruvate metabolism | dn,rf | pos |
| gs-00650 | Butanoate metabolism | dn | pos |
| gs-00640 | Propanoate metabolism | dn,gg,rf | pos |

## 4 Discussion

The most significant gene in the reference build, acyl-CoA diacylglycerol acyltransferase (*dgat*), plays a key role in the storage of intracellular triglycerides in eukaryotic organisms (Turchetto-Zolet et al., 2011). The only pathway that was significant in all transcriptome builds, propanoate metabolism (gs-00640), is also directly related to the storage of lipids. Propionic acid is the conjugate acid of propanoate and because propionic acid is one of the principal metabolites produced by gut microbiota (Al-Lahham et al., 2010), it can be hypothesized that the community structure of the gut microbiome plays a role in endurance metabolism. More in general from this work we observe that lipid metabolismis associated with the phenotypic extremes of endurance with evidence at the individual gene, functional module and pathway levels—although it is not a surprising find given that triglycerides, a derivative of glycerol, are among the most important molecules for energy storage in eukaryotic organisms (Athenstaedt and Daum, 2006).

Lipid metabolism may be a reoccurring theme, but the most surprising aspect of the results, when taken together, are the sheer number of diverse mechanisms at the disposal of individual frogs that appear to contribute to endurance heterogeneity. Take for example some of the most promising candidates among the 42 core genes. The glutamine transporter *slc38a3*, is a logical gene to contribute phenotypic heterogeneity, because the amino acid L-glutamine, like glucose is an essential substrate for many types of cells (Newsholme et al., 2003). Another core gene *nudt15* or *MTH2* in mice, has been shown to prevent mutations caused by reactive oxygen species, potentially placing the gene in a protective role (Cai et al., 2003). In humans, the gene *DAZAP1* is an associated with the RNA-binding protein *DAZ* suggesting a role at the level of transcription regulation (Tsui et al., 2000). The cyclic nucleotide degrading type 4 phosphodiesterase *pde4b* is an important upstream regulator involved in a wide variety of diseases like pulmonary fibrosis and B-cell lymphoma (Selige et al., 2011; Kim et al., 2011).

Functional annotations indicated that *btrc* is a modulator of a large variety of processes and pathways including some that involve NF-*κ*B and Wnt signaling (Wang et al., 2004). Appropriately, knockout mice for *baz1b* have a phenotype of ‘decreased circulating glucose level’ according to a phenotypic screen carried out by the International Mouse Phenotyping Consortium (Brown and Moore, 2012). The likely mechanism behind the observed phenotype is chromatin mediated regulation of transcription (Daxinger et al., 2013). Similarly, the top gene for the GG assembly, *ankrd17*, modulates gene expression through chromatin binding and its most relevant function is as a regulator of signaling pathways like NF-*κ*B (Deng et al., 2009; Wang et al., 2012). The functional coherence among the gene candidates, that potentially contribute to the observed distinction between endurant and non-endurant, is encouraging although it raises the question of how conserved the mechanisms are in vertebrates.

In humans, work has also been carried out to study the genetic differences between speed and endurance athletes and one of the most frequently cited markers of this distinction is the *α*-actinin-3 gene: *ACTN3* (*actn3* in *Xenopus*) (Yang et al., 2003). There has since been some focus on the R577X polymorphism of this gene the literature with respect to its association to various types of elite athletes. In this study none of the transcripts corresponding to *actn3* were significant, but the gene *act3* along with several other motor protein related genes are part of the 42 core genes. The gene *act3* (*ACTA1* in humans) is the predominant isoform of the filiment protein actin and nucleotide variants are associated with a number of myopathies (Laing et al., 2009). One could hypothesize that the compensatory mechanisms that come into effect with skeletal muscle deficiencies may play a role in the trade-off between endurance and speed. For specific interests, there is a certain appeal to identifying molecular markers of elite athletes and this work suggests a number of candidate genes and processes. Neither single nucleotide variants nor humans were within the scope of this study, but the homologs of the 42 core genes identified here could be scanned for variants in for example an exome sequencing case-control study.

The goal of this study was to elucidate genetics that help our understanding of how natural populations will respond and adapt to widespread environmental change as it is one of the greatest challenges facing ecologists today (Lawton and May, 1995; Balmford and Bond, 2005). In the last decade, predictive ecological models have become a popular tool for projecting the medium to long-term distribution and occurrence of species in response to environmental change (Raxworthy et al., 2003; Clark and Gelfand, 2006). Accurate predictions are, however, hampered by a wide range of assumptions and difficulties that include for example an efficient means to represent migration (Tylianakis et al., 2008). Precise ecological forecasting of population dynamics and persistence therefore requires a better integration of population movements and dispersal, as well as evolutionary potential in response to selection on mobility (Thuiller et al., 2008; Morales et al., 2010). Estimating the evolutionary trajectory of an organism will ultimately depend on our ability to identify the molecular underpinnings of variation in mobility. Another aspect of population dynamics that may need to be accounted for in future modeling efforts is the heterogeneity of behavioral syndromes, an important factor in some systems that has been largely overlooked (Cote et al., 2010). Finally, we focused on the skeletal muscle for the sake of specificity, but other tissues and organs like the brain play important roles in distinguishing endurance phenotypes (Yarrow et al., 2009).

RNA-Seq technologies are attractive to predictive modelers due to both the breath and depth of information contained within the sequencing results. From mapped reads it is possible to quantify differential expression, transcript variation (SNPs), alternative splicing (Wang and Cairns, 2013), and differential exon usage (Anders et al., 2012). Because a principal motivation for this study is to characterize and map gene function there was a focus on differential expression of transcripts. Further analyses of single nucleotide variants, alternative splicing and other transcriptome features would be helpful, particularly if the scope of the analyses were over a targeted phylogenetic spectrum with comparable phenotypes. Despite the lack of an evolutionarily perspective, the functional results presented here offer numerous avenues of further inquiry and the multi-level approach to expression analysis as applied over multiple transcriptome assemblies provides a natural way to prioritize the findings.

## 5 Methods

### 5.1 Experiments and Illumina transcriptome sequencing

*Xenopus tropicalis* were caught in the wild (December 2009) in Cameroon, brought back to France and housed at the Museum National d’Histoire Naturelle in Paris. Animals were placed in groups of 8-10 individuals in aquaria (60 × 30 × 30cm) at 24°C, which is assumed to be close to the preferred and optimal temperature of *Xenopus* frogs (see (Casterlin and Reynolds, 1980; Miller, 1982)) and is similar to water temperatures measured in the field (22–26°C). Frogs were fed every other day with beef heart, earthworms or mosquito larvae *ad libitum*. All individuals were given one month to recover and were then pit tagged (Nonatec, Rodange, Luxembourg). We measured endurance on a subset of 8 males through performance tests as in (Herrel and Bonneaud, 2012). Individuals were sacrificed via an intramuscular overdose of Imalgene (ketamine hydrochloride, 100mg/ml) injected in the left leg. This allows a profound anesthesia eventually resulting in the death of the animal. While under deep anesthesia, the knee extensor muscles of the right leg were extracted for subsequent RNA sequencing.

Tissues were extracted, transferred to labeled tubes containing RNA-later and conserved at −80°C until further processing. Muscle samples were subsequently washed in RNAse free ultra pure water and muscle cells were destroyed using a Trizol solution; samples were then mixed for 1 to 2 min using an Ultra-Turrax (IKA Inc. at Staufen, Germany). All equipment was washed between each sample with a specific wash protocol (SDS 1%, Ethanol 70%, Rnase Zap and water). Samples were then centrifuged for 10 minutes at 4°C and 12000g and supernatant was transferred to a Phase-Lock-Gel and incubated for 5 min at room temperature. Chloroform was added (0.2ml per ml of Trizol) and samples centrifugated for 20 min at 4°C and 2500g. Isopropanol (0.5 ml per ml of trizol) was then added to supernatants and incubated overnight at −20°C. Samples were centrifugated for 10 minutes at 4°C and 12000g and pellets were washed in ethanol 70% (1ml per ml of Trizol). Again the samples were centrifuged at 7500g for 5 min and then dried on ice. Next the samples were then dissolved in RNAse free ultra pure water and stored at −20°C. RNA concentration and quality were verified on a Nanodrop (Thermo Fisher Scientific at Waltham, Massachusetts, USA) and an Agilent 2100 BioAnalyser (Agilent technologies at Santa Clara, California, USA). The samples were hybridized using a single lane on an Illumina HiSeq 2500 machine located at the University of Exeter Sequencing Service core facility. Phasing corrections were made against the standard PhiX control lane.

### 5.2 Differential analysis pipeline

Reads were trimmed and adapter sequences were removed using Trimmomatic (Bolger et al., 2014) with the following parameters: LEADING:5 TRAILING:5 SLIDINGWINDOW:4:15 MINLEN:36. The reads were *de novo* assembled both with and without a reference genome (genome-guided) using Trinity (Grabherr et al., 2011). For both of the Trinity assemblies, trimmed reads were aligned back to the assembly with Bowtie2 (Langmead and Salzberg, 2012) and abundance was estimated using RSEM (Li and Dewey, 2011). The most recent stable build of the *X. tropicalis* genome (downloaded from Xenbase (Bowes et al., 2008; James-Zorn et al., 2013)) (Version 7.1) was used for the genome-guided and reference assemblies. For the reference assembly, reads were mapped to the main scaffolds using STAR Dobin et al. (2013) and the annotation file provided by Xenbase. The resulting SAM files were converted for use with downstream tools with SAMtools (Li et al., 2009). The reference genome alignments were used to determine gene-level counts and to estimate coverage by making use of the Python package HTSeq Anders et al. (2015).

The analysis pipeline is generalized as a flow chart in Figure 5. The three methods for transcriptome assembly are shown in the context of the overall differential expression analysis framework. The three routes for arriving at identified genes of interest are indicated by arrows with different shades gray and black. This study focused on expression characterized by read counts, but there are other features like splicing events, exon usage, and SNVs that can be equally studied using a variation of this framework. The individual transcript level analyses were performed with the R package DESeq2 (Love et al., 2014). The GSA portion of the pipeline contains both the functional module and pathway analyses. A recent publication details the method and rationale for producing functional modules (Richards et al., 2015). Both the functional modules as well as the pathways downloaded from KEGG were tested for significance using the R package GSA (Efron and Tibshirani, 2007).

**Figure 5:**
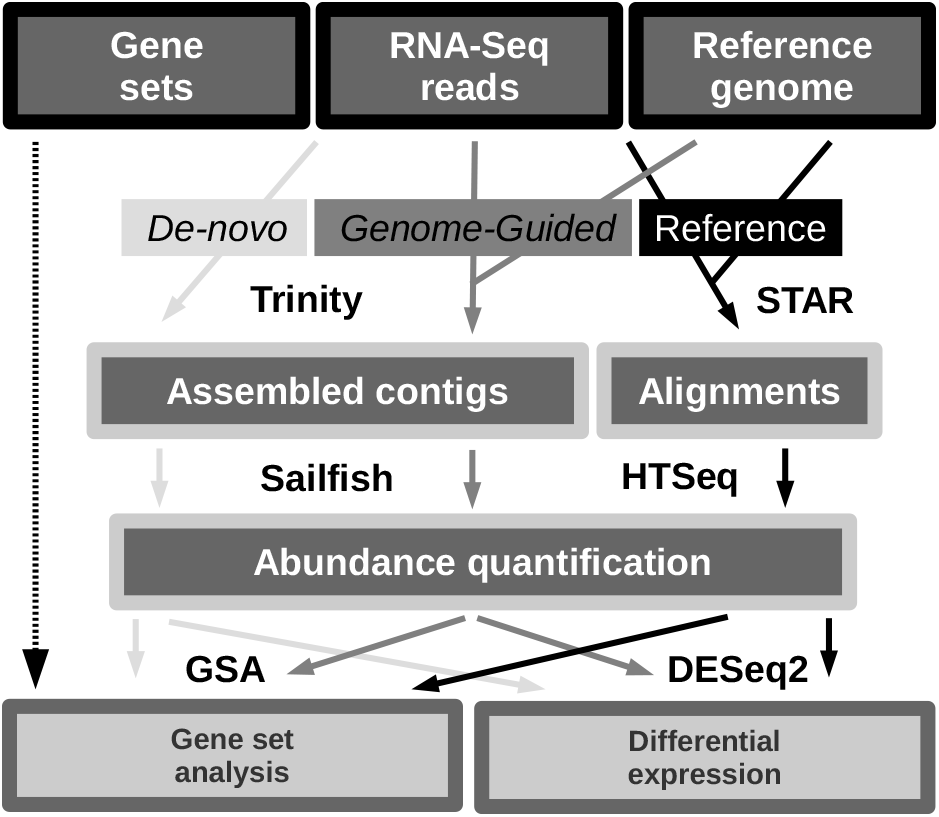
Generalized pipeline. The conceptual diagram begins with gene sets that are determined before the analysis, reads that have been quality controlled and a reference genome. Transcriptomes are then assembled as contigs or via alignments with a reference genome. The *de novo* and genome guided methods are both technically *de novo* builds, but the genome guided make use of a reference genome. The reference build is created by aligning reads. Gene and isoform level counts along with other genomic features are then extracted from the builds. Finally, individual gene and gene set level significance analyses are undertaken. The arrow with different shades indicate the three distinct assemblies and the unboxed text corresponds to the software used.

*X. tropicalis* build sequences were BLASTed (Camacho et al., 2009) against the SwissProt portion of the UniProt database (Consortium, 2014) to enable mapping of identifiers to gene sets derived from multiple taxa (e-value threshold 1*e* – 05). The gene set generation, BLAST, namespace mapping and plotting was carried out using the Python library htsint (Richards et al., 2015). The Python packages NetworkX (Hagberg et al., 2008) and Matplotlib Hunter (2007) were used for algorithms that are implemented as part of the library. All analysis methods, including code snippets are available as part of the Supplemental methods.

### 5.3 Gene set analyses

In this study we report the results in terms of NCBI gene identifiers due to stability, but also because the gene-namespace is smaller than the protein-namespace which reduces the compute time required to partition the transcriptome into gene sets. The functional modules were created as detailed in Richards et al. (2015) using the *biological Process* and *molecular function* aspects of the Gene Ontology. Using hstint’s underlying database we retrieved all of the gene and term mappings that were specific to *Xenopus laevis* and *Xenopus tropicalis* and using spectral clustering the functional modules were created by partitioning all known *Xenopus* genes based on gene-gene functional similarity. The functional similarity was quantified through a graph-based method that integrates the annotation data from both frog species.

The original work by Mootha et al. (2003) and Subramanian et al. (2005) proposed a straightforward procedure where first the individual genes in a set are ranked based on magnitudes of their differential expression with respect to the outcome variable. Then an enrichment score, that is an agglomeration of individual scores, is calculated and finally statistical significance was assessed using a permutation approach. Efron and Tibshirani (2007) improved this method by proposing the use of a *maxmean* test statistic, in place of a weighted Kolmogorov Smirnov test. GSA has become the standard way to investigate predefined gene lists, like biological pathways (Tian et al., 2005), that come from a variety of high-throughput technologies (Huang et al., 2009; Hung et al., 2012).

## 6 Conclusions

It is now widely accepted that human activities are the primary driver of the rapid worldwide decline of species currently taking place, with factors like habitat destruction playing an important role. Population forecasting efforts are hampered by an inability to accurately model dispersal and mobility. Dispersal and animal movement in general is complicated by phenotypic heterogeneity as well as other factors like epistasis and genotypic heterogeneity. Some of the genetic elements that underlie endurance capacity can be generalized to pathways or functional modules, but there remain individual genes that are potentially causal by themselves (i.e. *act3*). Further experimentation is required to tease out the relationships among the important individual genes and to determine the extent of exchangeability among member of a functional module or pathway. Specifically, the role of lipid storage and lipid metabolism has presented itself as a likely suspect in explaining endurance heterogeneity in frogs. The multi-level approach to gene expression analysis is readily generalizable to other species and the use of multiple transcriptome assemblies to help prioritize and navigate results will likely be most useful in non-model organisms that are poorly annotated, but given that *de novo* assembly methods are improving their usefulness may eventually be proven for even the most widely studied organisms.

## 7 Data access

The transcriptome data from this study have been submitted to the NCBI BioProject (https://www.ncbi.nlm.nih.gov/bioproject) (XXXXXX) and to the NCBI Gene Expression Omnibus (https://www.ncbi.nlm.nih.gov/geo) using the accession number GSEXXXXXX.

## Acknowledgements

### 8 Acknowledgements

We would like to thank Michel Baguette and Hervé Philippe for providing valuable resources and discussion. Support for this work was provided by the Agence Nationale de la Recherche (ANR; France) MOBIGEN [ANR- 09-PEXT-003].

